# *Francisella novicida* Cas9 interrogates genomic DNA with very high specificity and can be used for mammalian genome editing

**DOI:** 10.1101/591263

**Authors:** Sundaram Acharya, Arpit Mishra, Deepanjan Paul, Asgar Hussain Ansari, Mohd. Azhar, Manoj Kumar, Riya Rauthan, Namrata Sharma, Meghali Aich, Dipanjali Sinha, Saumya Sharma, Shivani Jain, Arjun Ray, Suman Jain, Sivaprakash Ramalingam, Souvik Maiti, Debojyoti Chakraborty

## Abstract

Genome editing using the CRISPR Cas9 system has been used to manipulate eukaryotic DNA and make precise heritable changes. Although the widely used *Streptococcus pyogenes* Cas9 (SpCas9) and its engineered variants have been efficiently harnessed for numerous gene-editing applications across different platforms, concerns remain, regarding their putative off targeting at multiple loci across the genome. Here we report that *Francisella novicida* Cas9 (FnCas9) shows a very high specificity of binding to its intended targets and negligible binding to off-target loci. The specificity is determined by its minimal binding affinity with DNA when mismatches to the target sgRNA are present in the sgRNA:DNA heteroduplex. FnCas9 produces staggered cleavage, higher HDR rates and very low non-specific genome editing compared to SpCas9. We demonstrate FnCas9 mediated correction of the sickle cell mutation in patient derived iPSCs and propose that it can be used for precise therapeutic genome editing for a wide variety of genetic disorders.

**SIGNIFICANCE STATEMENT:** Therapeutic genome editing has been significantly accentuated by the advent of CRISPR based gene correction. However, genomic off-targeting has been a major setback for clinical translation. Although high fidelity versions of Cas9 have been rationally designed, they recognize and bind to off-targets. In this study, we characterize a naturally occurring Cas9 from *Francisella novicida* (FnCas9) that shows negligible binding affinity to off targets differing by one or more mismatches, rendering it highly specific in target recognition and editing. We show that FnCas9 can direct both HDR and NHEJ mediated DNA repair, generates higher rate of HDR and negligible off-target editing. Finally we show its potential in therapeutic genome editing by correcting the sickle cell anemia mutation in patient derived iPSCs.

## INTRODUCTION

The process of introducing changes in the DNA of cells using Clustered regularly interspersed short palindromic repeats (CRISPR)-CRISPR-associated (Cas) proteins has emerged as a powerful technique in molecular biology with potentially far reaching applications in gene therapy (1–6). The method involves harnessing the prokaryotic type II CRISPR-Cas protein Cas9, which in complex with a single guide RNA (sgRNA), can be directed to any region in the DNA upstream of a protospacer adjacent motif (PAM) with which the sgRNA sequence finds a match (7–9). Upon double stranded cleavage at the target site, the endogenous repair machinery of the cell can be utilized to make nucleotide changes in the DNA (6,10–12). Although several Cas9 proteins recognizing different PAM sequences have so far been reported in literature, only a subset of these have been characterized and have demonstrated genome-editing ability in eukaryotic cells (5,13–16).

Cas9 from *Francisella novicida* (FnCas9) is one of the largest Cas9 orthologs and has been shown to predominantly interact with the 5’-NGG-3’ PAM motif in DNA (17). The crystal structure of FnCas9 ribonucleoprotein (RNP) in complex with target DNA has revealed both conserved and divergent features of interaction that is unique among the Cas9 enzymes studied. Unlike SpCas9, FnCas9 does not form a bilobed structure, has a different sgRNA scaffold and has been implicated in RNA targeting (17–18). By structure guided protein engineering, FnCas9 can be made to recognize a 5’-YG-3’ PAM (17). Although the protein can efficiently cleave DNA substrates *in vitro*, its *in vivo* activity at several genomic loci is significantly diminished as compared to SpCas9 (17,19).

FnCas9 has been reported to have high intrinsic specificity for its targets, most notably by tolerating only a single mismatch in the sgRNA at the 5’ position of the PAM distal end (17). This is in stark contrast with SpCas9, which has shown variable levels of off targeting due to tolerance of mismatches predominantly in the ‘non-seed’ region in the sgRNA, wherever these are encountered in the genome (20). To what extent FnCas9 mediates this high specificity of target interrogation is not known and whether these properties can be harnessed for highly specific genome editing at a given DNA loci has not been investigated. The distinct structural attributes of FnCas9 and its low tolerance for mismatches led us to investigate its DNA interaction properties and role in genome editing.

## RESULTS

### FnCas9 shows sequence dependent DNA binding affinity and cleavage kinetics

The interaction of Cas9 to its substrate occurs in a multi-step process involving PAM binding, target unwinding and DNA:RNA heteroduplex formation (21–22). As several components in this process are dependent on enzyme conformation, we speculated that FnCas9 might possess DNA interrogation parameters that are intrinsically linked to its structure. To dissect the binding affinity of FnCas9 RNP complex with target DNA, we used MicroScale Thermophoresis (MST) to determine the dissociation constant (K_d_) of FnCas9:target interaction. To perform binding affinity measurements, we generated recombinant catalytically *dead* (inactive) FnCas9 (dFnCas9) tagged with green fluorescent protein (GFP) and confirmed its inability to perform *in vitro* cleavage of a DNA substrate (SI Appendix, Fig. S1A,B). Electrophoretic gel mobility shift experiments (EMSA) concluded that FnCas9 and its sgRNA sequence interaction reaches saturation at a molar ratio of approximately 1:1.5, similar to that reported for SpCas9 binding with its sgRNA (9,23; SI Appendix, Fig. S1C).

Next, we investigated the binding of dFnCas9: sgRNA RNP complex with two different DNA substrates (*VEGFA3* and *EMX1*) and observed a K_d_ of 150.7 ± 36.6 nM and 78.8 ± 18.5 nM respectively (Fig. 1). To compare the binding of dSpCas9 to the same targets, we purified recombinant dSpCas9-GFP and interrogated the substrates using MST. We observed lower K_d_ of substrate binding for dSpCas9 for each of the targets (49.6 nM ± 9.78 nM and 10.9 ± 6.2 nM respectively) suggesting that in comparison with dFnCas9, dSpCas9 might have a generally higher affinity for the same substrate sequence (Fig. 1). Notably, K_d_ of SpCas9 binding to its substrates as reported in literature using other techniques (such as EMSA, Beacon assays or Active site titration assays) also fall in a similar range (≤ 100nM) as observed under our experimental conditions, suggesting concordance between the different methods used in determining binding affinity (9,24–27). We tested two more substrate sequences (*EMX1_2, c-MYC*) with dFnCas9 and once again observed a wide distribution of dissociation constants (K_d_ 462 ± 52.4 nM and 30.6 + 6.7 nM respectively, SI Appendix, Fig. S1D). Collectively, this suggests that FnCas9 shows a diverse range of binding affinities with different DNA substrates and its affinity for targets is generally lower than that of SpCas9. We then proceeded to investigate if these differences in binding affinity might lead to variability in cleavage efficiencies across different substrates. To investigate substrate cleavage, we purified wild type FnCas9 and SpCas9 and performed *in vitro* cleavage (IVC) assays. When different DNA substrates were incubated with equal amounts of FnCas9 RNP, we observed that cleavage reached completion at different time points (30 min – 2h) for these substrates, suggesting that the sequence of the target might affect the rate of cleavage; SpCas9 however, did not show such variability with any of the targets (Fig. 2). Interestingly, when we took a substrate that showed gradual completion of cleavage (*c-MYC*) and performed IVC with increasing molar concentrations of RNP, we observed that substrate cleavage completion could be shifted to an earlier time-point upon incubation with higher concentration of FnCas9 RNP (SI Appendix, Fig. S2A). Thus, FnCas9 cleavage efficiency varies proportionally to RNP concentration indicating that it might act as a single turnover enzyme, a feature reported earlier for SpCas9 (9). Collectively, these results suggest that FnCas9 has a generally high threshold for substrate recognition and even in the case of complete match of crRNA with its target, exhibits different cleavage kinetics when encountering different target sequences.

**Figure 1.**
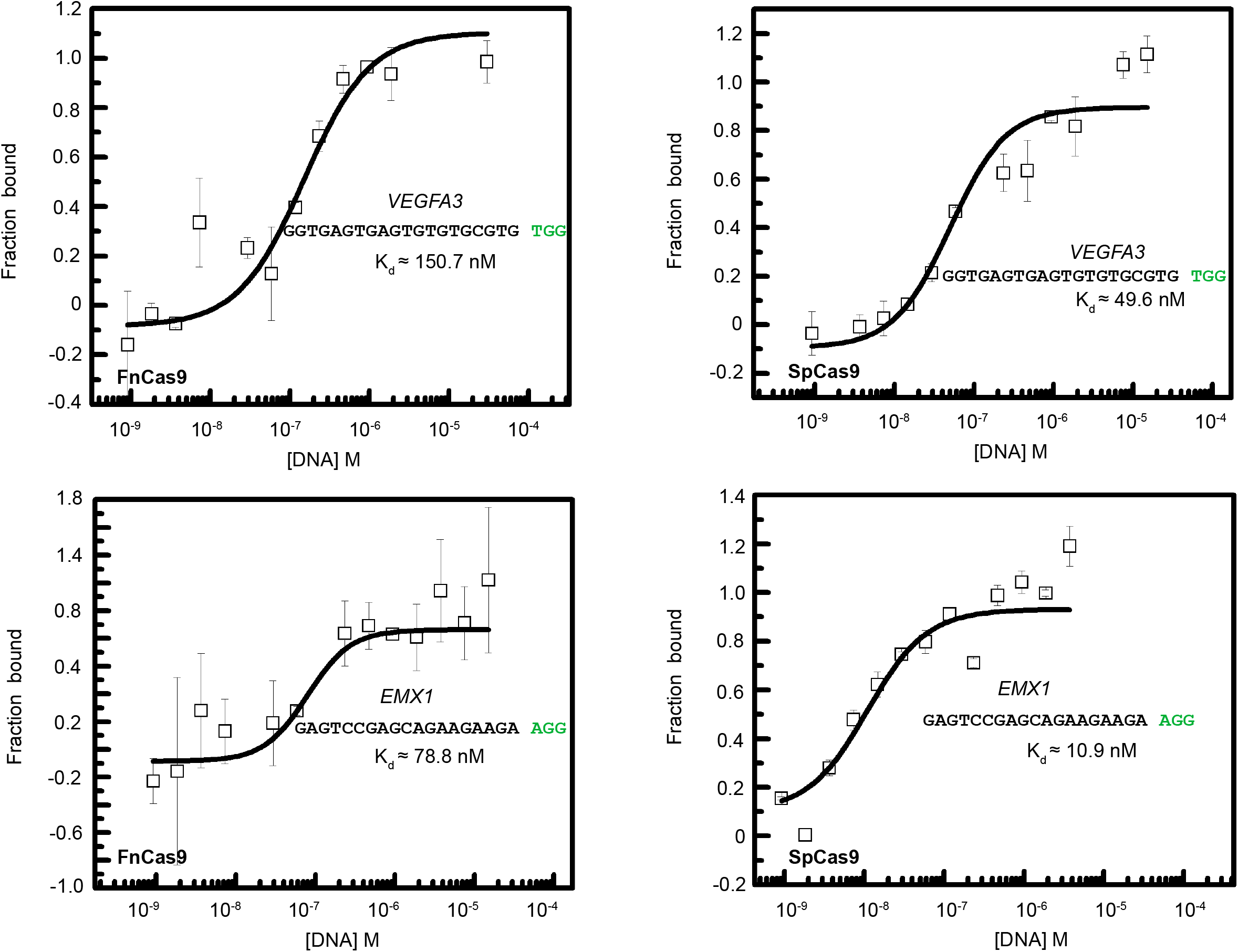
FnCas9 shows different affinity and cleavage kinetics for substrates as compared to SpCas9. MST results showing binding of dFnCas9-GFP (left) and dSpCas9-GFP (right) to two substrates, *VEGFA3* (top) and *EMX1* (bottom) expressed as normalized fluorescence units (y-axis) with respect to varying concentrations of purified substrate (x-axis). The substrate sequences are indicated in the box with the PAM shown in Green. Error bars represent SEM (2 independent experiments).

**Figure 2.**
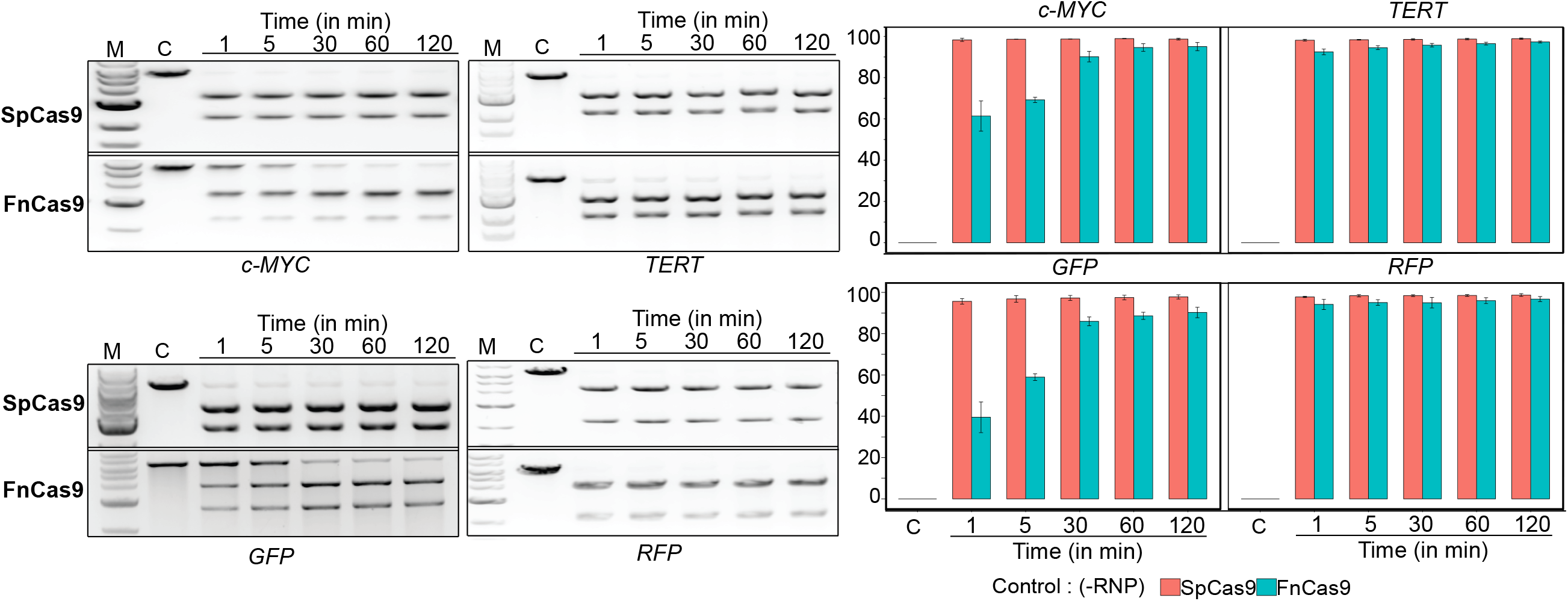
FnCas9 shows different cleavage kinetics for substrates as compared to SpCas9. *In vitro* cleavage assay showing activity of WT SpCas9 and WT FnCas9 (50nM Cas9:150nM sgRNA) on indicated substrates (linearized plasmids, 250ng) at different time points (1 min – 2h). Top band represent uncleaved target while the two bottom bands represent cleaved products. Quantification is shown on the right, y-axis represent percentage cleavage. Error bars represent SD (3 independent experiments, C = control).

### More than one mismatch in sgRNA:DNA heteroduplex abolishes FnCas9 cleavage

The specificity of a genome editing protein is guided by a balance between its affinity for target and ability to discriminate off targets. A recent report had suggested that FnCas9 shows higher intrinsic specificity than SpCas9 to its target by showing less tolerance to single mismatches at certain sgRNA positions (19). We investigated the *in vitro* cleavage efficiency of both SpCas9 and FnCas9 by systematically changing every base in a given substrate and observed that whereas SpCas9 cleaved nearly equally at all mismatched positions, FnCas9 was less tolerant to single mismatches and was particularly stringent at base positions 19, 18, 17 and 16 at the PAM distal end (Figure 3) where considerable abolishment of cleavage was observed (~ 56 ± 7%). Interestingly, engineered highly specific Cas9 variants that are able to prevent cleavage at off targets containing PAM distal mismatches adopt a conformational structure that renders the HNH DNA cleavage domain inactive (24). Recent biophysical studies using single molecule fluorescence resonance energy transfer (smFRET) have revealed a highly dynamic conformation of SpCas9 where allosteric interactions between the PAM distal end and the HNH domain of the Cas9 enzyme renders a cleavage-impaired conformationally ‘closed’ configuration upon encountering mismatches close to the 5’ end of the sgRNA (28–29). To dissect if a greater stringency of target recognition could be achieved in case of FnCas9 by increasing the number of mismatches at the 5’ end of the substrate, we selected two well-studied loci *EMX1* and *VEGFA3*, amplified their genomic off-targets with 2 and 3 mismatches at the non-seed region (PAM distal end) and interrogated the *in vitro* cleavage efficiency of FnCas9. Remarkably, FnCas9 was unable to cleave the substrate in the presence of 2 or 3 mismatches for both loci suggesting that it is extremely specific in target recognition, particularly when the mismatches occur together in the PAM distal region (Fig. 4A). SpCas9 however, was able to cleave both the mismatched substrates (Fig. 4A). Previous studies have highlighted the importance of defined mutations in the REC3 domain of highly specific engineered versions of Cas9 in determining target specificity by allosterically regulating the HNH domain from adopting a cleavage-competent form (24,30). However, the different engineered variants showed similar binding affinity for their off-targets as wild type SpCas9, even though they did not cleave these targets suggesting that they probably remain bound to off-targets in a cleavage incompetent state (24). We asked if FnCas9 too shows similar properties and investigated the binding affinity for off-targets using MST. Strikingly, FnCas9 showed negligible to no binding affinity for substrates having two mismatches where no cleavage was observed suggesting that it either interacts extremely weakly or is evicted from the substrate following off-target interrogation (Fig. 4B, SI Appendix, Fig. S2A). SpCas9 however, showed a strong binding affinity for both on- and off-targets as reported previously (24) (Fig. 4B, SI Appendix, Fig. S2B). Taken together these results indicate that FnCas9 has a fundamentally distinct outcome of off-target recognition and binding as compared to SpCas9 and its engineered derivatives.

**Figure 3.**
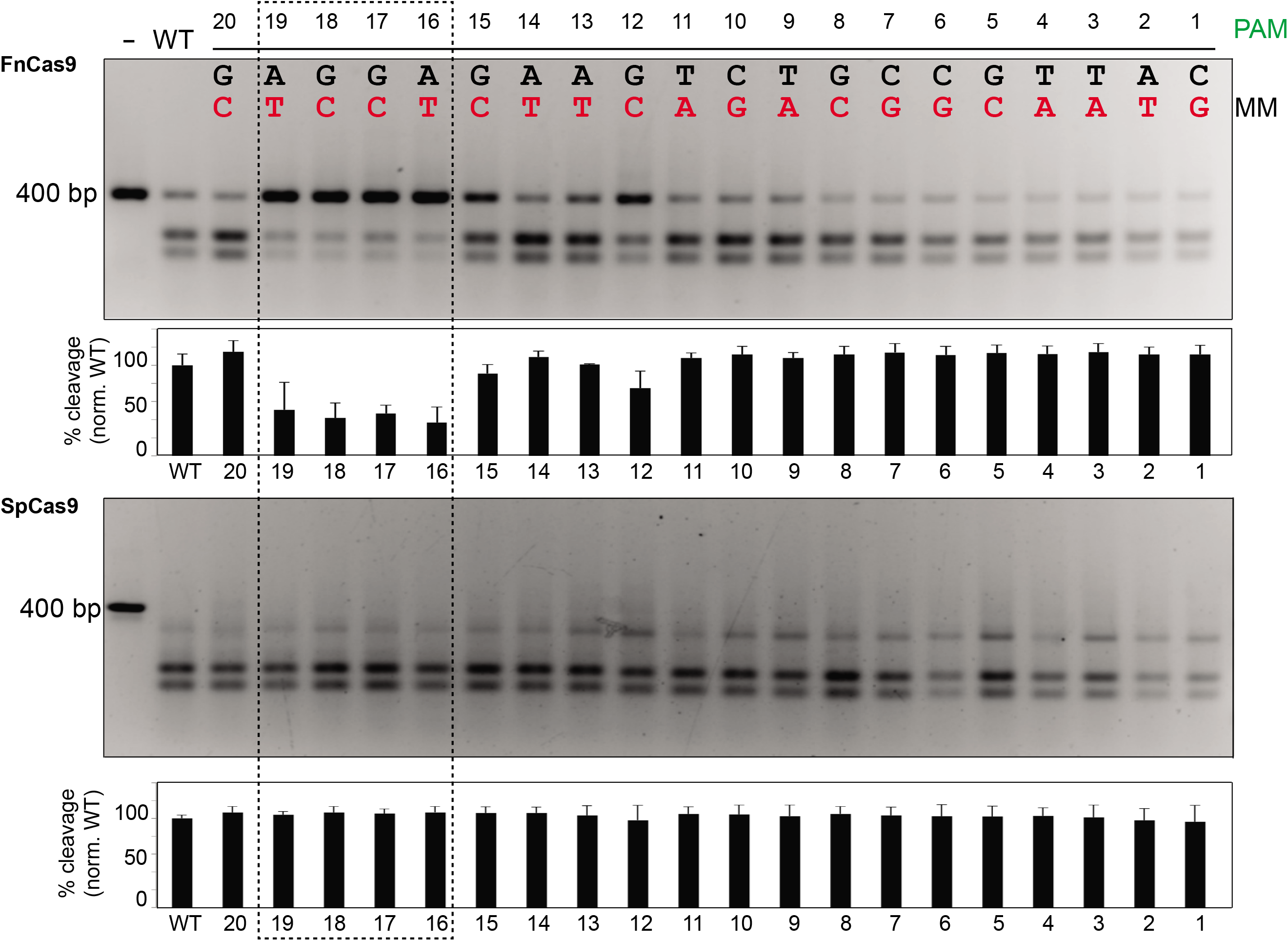
Mismatches in target abrogates FnCas9 cleavage activity. Mismatches at different positions along the substrate alter FnCas9 cleavage outcomes while retaining SpCas9 activity. Representative *in vitro* cleavage outcome of HBB substrate (purified PCR product, 100ng) showing mismatches at a single position in every lane (indicated in red, other bases remaining unchanged) interrogated with FnCas9 (top) or SpCas9 (bottom) in the form of RNP complexes (250nM) is shown, PAM is indicated in green. Top band represents uncleaved target while the two bottom bands represent cleaved products. Quantification for each reaction is shown below the gel images. Error bars represent SD (3 independent experiments).

**Figure 4.**
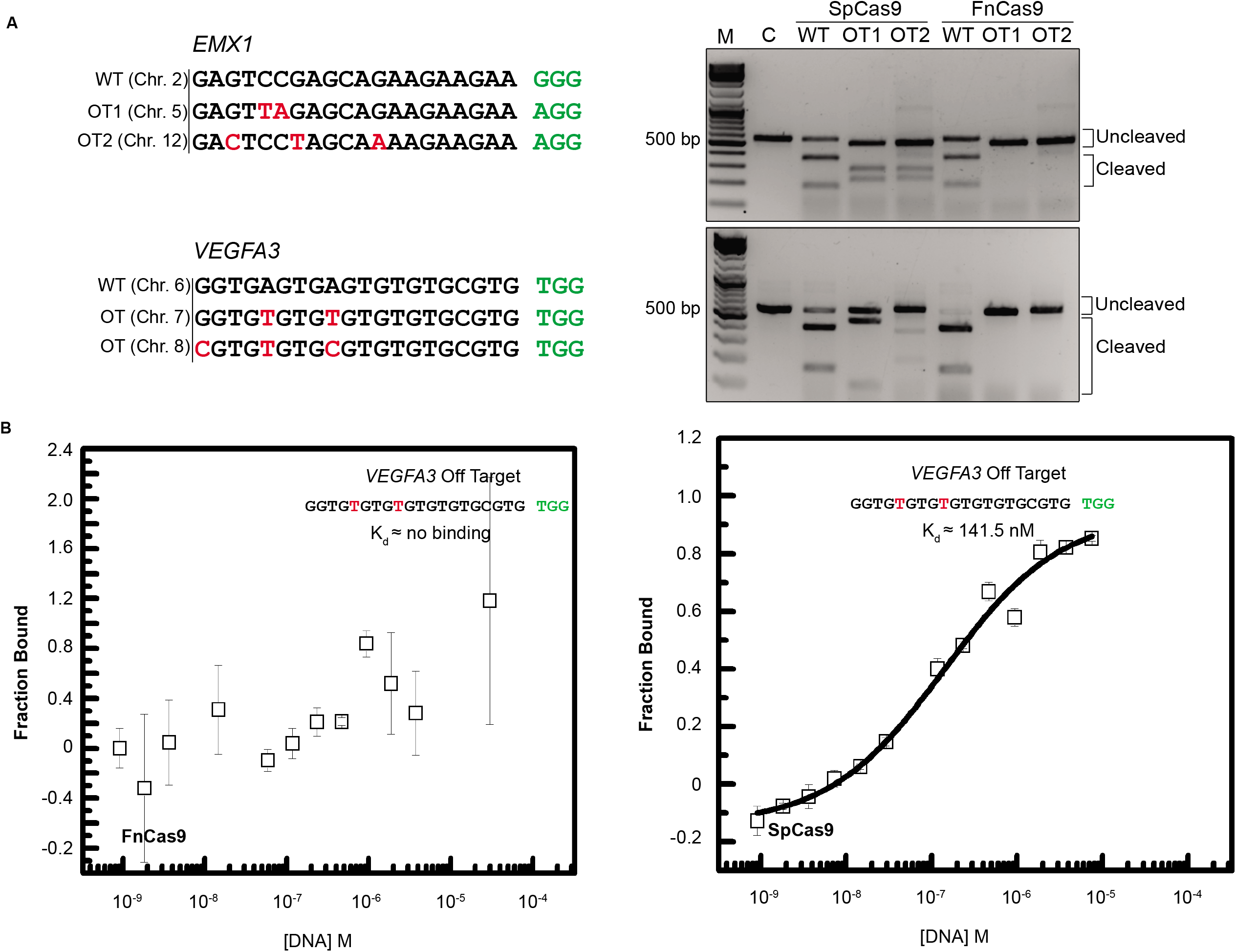
FnCas9 does not bind to mismatched substrates. (A) FnCas9 does not tolerate 2 or more mismatches in the substrate at the PAM distal end. *In vitro* cleavage outcomes of *EMX1* (top) and *VEGFA3* (bottom) targets interrogated by WT FnCas9 and SpCas9 are shown (right). On the left, the sequences with 2 or more mismatches (indicated as red bases) are presented. PAM is indicated in green. Data representative of 3 independent experiments. (B) MST results showing binding outcomes of dFnCas9-GFP (left) and dSpCas9-GFP (right) to *VEGFA3* off-target substrate expressed as normalized fluorescence units (y-axis) with respect to varying concentrations of purified substrate (x-axis). The substrate sequences are indicated in the box with the PAM shown in Green. Error bars represent SEM (2 independent experiments).

Intrigued by the ability of FnCas9 to discern mismatches in the target with high accuracy, we looked at the protein structure to understand the basis for this specificity. The interaction of FnCas9 with its substrate is predominantly mediated by an expanded REC3 and the associated REC2 domains that are structurally distinct from those found in SpCas9 or *Staphylococcus aureus* Cas9 (SaCas9) (17). In the crystal structure of FnCas9, the REC3 domain was reported to contain a structural zinc ion in a coordination sphere consisting of 4 cysteine residues (17). Notably, complete abolishment of FnCas9 activity was observed when *in vitro* cleavage reaction was supplanted with Zn^2+^ (1mM) suggesting that the binding of the metal ion inhibits FnCas9 mediated cleavage even in the presence of Mg^2+^ (SI Appendix, Fig. S2C). Upon analyzing the crystal structure of FnCas9 RNP in complex with target DNA, we observed that the FnCas9 has a much higher electrostatic potential than SpCas9 and interacts with several bases both on the PAM distal and PAM proximal ends of the substrate (SI Appendix, Fig. S2D). We speculate that these interactions might be necessary for FnCas9 to adopt a cleavage competent state and in the presence of at least 2 mismatches, the complex dissociates. In fact, the electrostatic contributions of FnCas9 on each of the 20 bases in the substrate are higher than that of SpCas9, possibly leading to stringent interrogation and subsequent dissociation upon mismatch recognition (SI Appendix, Fig. S2E). We asked if these features could make FnCas9 extend its specificity for off-target discrimination at both the PAM proximal and distal ends. To test this, we designed substrates carrying the therapeutically relevant human *HBB* sequence and introduced two mutations at different positions throughout the substrate. We observed that cleavage activity went down drastically, suggesting that FnCas9 has very low tolerance for double mismatches along most of the sgRNA:DNA duplex, while SpCas9 activity remained largely unaffected (SI Appendix, Fig. S3A). This discrimination of off-targets is not correlated with distance between the two mismatches (SI Appendix, Fig. S3A). We also inquired if off-targets can be distinguished by introducing double mismatches in the sgRNA sequence. To this end, we targeted this substrate using sgRNAs having double mismatches at various positions in the PAM proximal and distal ends and observed that complete loss of cleavage activity could be effected with mismatch combinations both at PAM proximal and distal ends. (SI Appendix, Fig. S3B). To rule out that this specificity is dependent on the sequence being interrogated, we looked at three more targets (*c-MYC, VEGFA3 and EMX1*) with sgRNAs containing 2 mismatches and found that in these targets too, cleavage was completely abrogated in the presence of these mismatches (SI Appendix, Fig. S3C). Taken together these results suggest that the trigger for mismatch discrimination is embedded in the FnCas9 structure and its mode of target interrogation and not the DNA sequence that it engages with.

### FnCas9 mediates cellular genome editing with very high precision

Intrigued by the high specificity of substrate recognition under *in vitro* conditions, we next investigated if FnCas9 can function *in vivo* as a genome-editing agent and how its genome editing properties compared with that of SpCas9. We first investigated if FnCas9 can access nuclear DNA by complexing dFnCas9-GFP protein with sgRNAs that target the Telomeric TTAGGA repeats and performing in situ labeling of genomic loci in mouse ES cells that are known to have longer telomeres amenable to imaging (31). We observed identical nuclear punctate dots for both dSpCas9-GFP and dFnCas9-GFP using deconvolution microscopy establishing that FnCas9 can localize to genomic DNA efficiently (Fig. 5A). We then asked if DNA localization also translates to actual DNA binding events. To this end, we used a mouse cell line containing GFP sequence (Neuro2A-GFP), transfected it with a plasmid expressing FnCas9 and *in vitro* transcribed sgRNAs targeting the GFP sequence and measured chromatin binding of FnCas9 using Chromatin immuno-precipitation (ChIP). We observed up to 100-fold enrichment of the protein at the target after 24 hours post sgRNA transfection (SI Appendix, Fig. S3D) thus confirming that both active and dead FnCas9 were able to access their target sites on DNA in two different mouse cell lines.

**Figure 5.**
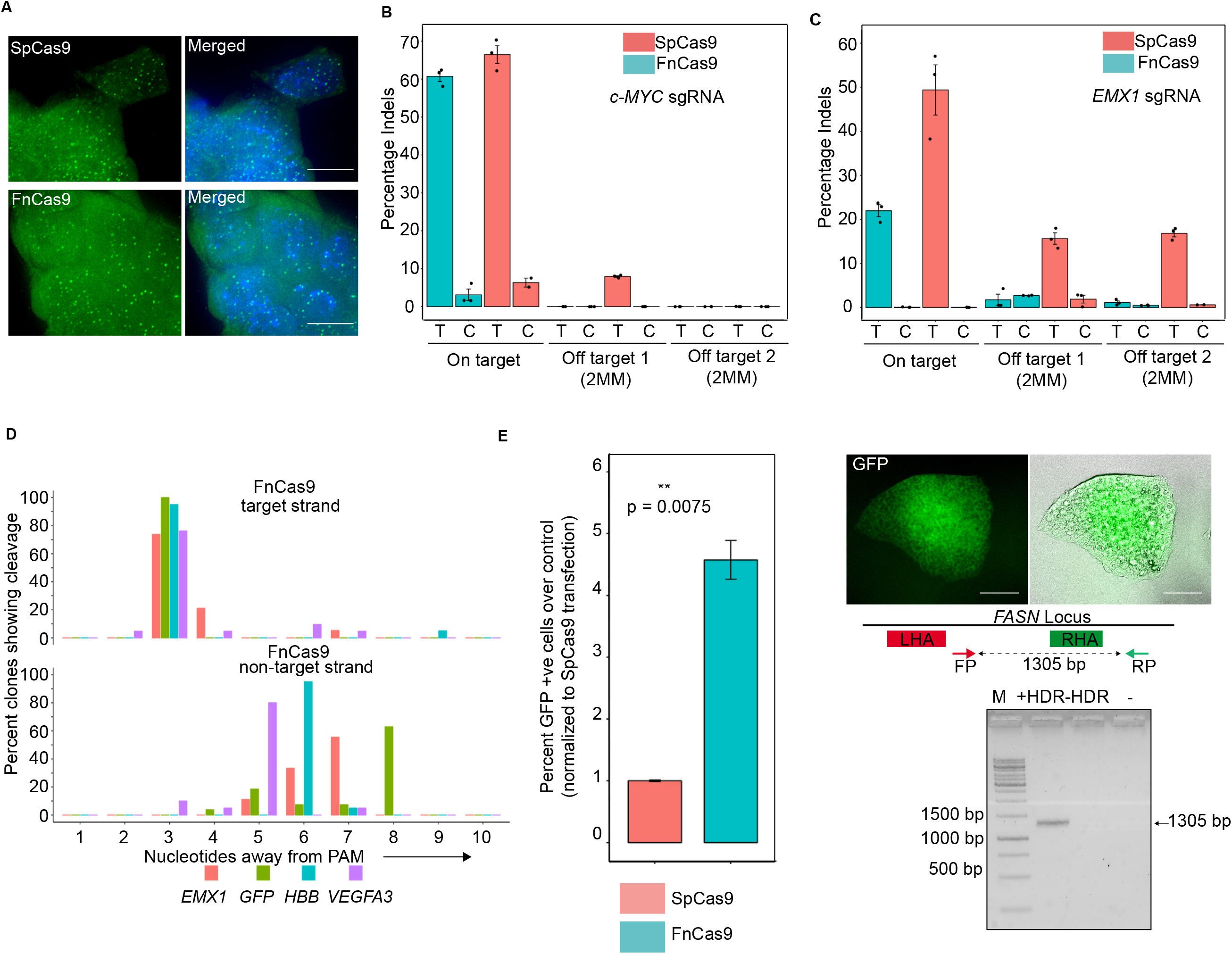
FnCas9 performs genome editing with very high specificity. (A) Representative Immunofluorescence image showing the nuclear localization of SpCas9-NLS-GFP or FnCas9-NLS-GFP protein (shown in green) with sgRNA (as RNP complex) against telomeric repeats in mouse ES cells. DNA is counterstained with DAPI and represented in blue. Scale bar 10 μm. (B) Indel events (expressed as percentage) obtained by amplicon sequencing upon SpCas9 or FnCas9 targeting *EMX1* in HEK293T cells. Individual data points from independent experiments are shown. Error bars represent SEM (3 independent experiments except for on target control for SpCas9). T = Test, C = Control. (C) Indel events (expressed as percentage) obtained by amplicon sequencing upon SpCas9 or FnCas9 targeting *c-MYC* in HEK293T cells. Individual data points from independent experiments are shown. Error bars represent SEM (3 independent experiments). T = Test, C = Control. (D) Cleavage positions on target and non-target strand by FnCas9 determined using Sanger sequencing for different targets indicated. Y axis represents percentage of sequenced clones showing cleavage at a given nucleotide. X axis represents the position of the base away from PAM in the sgRNA. (E) HDR is represented as percentage of GFP positive cells normalized to SpCas9. Student’s t-test p-values are shown. A representative image of GFP tagged HEK293T cells after FnCas9 mediated HDR is shown, scale bar 50um. Genotyping PCR confirming targeted GFP integration at the FASN locus in sorted GFP positive (+HDR) or negative (-HDR) and no template control (-) for FnCas9 mediated HDR. Integration at desired locus generates a 1305bp PCR product.

Having established that FnCas9 localizes and binds to DNA in mouse cells, we next asked if it can perform genome editing in human cells and how this compares with SpCas9. For a bona-fide comparison between the two different Cas9 proteins with identical expression parameters, we generated constructs with either SpCas9 or FnCas9 and their corresponding sgRNAs from the same backbone. To select cells that received this plasmid by FACS sorting, we supplemented this construct with a T2A-eGFP sequence. To compare on-target and off-target editing, we selected 2 loci *c-MYC* and *EMX1* where genomic off-targets with 2 mismatches could be identified and 1 loci *HBB* where genomic off-targets with 3 mismatches could be identified. In the case of *EMX1*, these off-targets were earlier validated by an unbiased genome-wide GUIDE-Seq study (32). We performed genome editing with SpCas9 and FnCas9 in HEK293T cells and a construct containing a scrambled sgRNA sequence was used as control. After sorting GFP positive cells from both SpCas9 and FnCas9, we isolated genomic DNA from these cells, amplified the target and off-target loci for each of the genes and performed deep sequencing to quantify the number of insertions/deletions (indels). We observed that FnCas9 was able to cleave all the 3 loci efficiently although with different efficiencies as compared to SpCas9. In the case of *c-MYC*, ~60% indels were seen which dropped to ~22% for *EMX1* and ~19 % for *HBB*. In contrast, SpCas9 showed greater than 50% indels for each of the 3 loci tested (Fig. 5B-C, SI Appendix, Fig.S3E). Analysis of the percentage of either insertions or deletions at the different loci did not reveal a consistent trend for any of these events being favored both SpCas9 or FnCas9 (SI Appendix, Fig. S4A).

Although SpCas9 cleaved more efficiently at each of the 3 loci, it showed modest to high cleavage at the off-targets for each loci. Among those with 2 mismatches, these included one off target for *c-MYC* (~8%) and 2 off targets for *EMX1* (~16% each, Fig. 5B-C). Even with 3 mismatches to the sgRNA it showed ~2% cleavage for at least 1 off-target. Strikingly, FnCas9 did not show any cleavage at any of the off-targets tested except *EMX1* off-target 2, where a very low ~1.15% indels were detected (Fig. 5C). Of note, the off-targets previously validated for cleavage by SpCas9 through GUIDE-Seq (*EMX1*) also showed the highest SpCas9 off-target activity in our analysis suggesting concordance between the two studies in terms of bona-fide off-targeting. FnCas9 however, showed almost no cleavage at these sites either suggesting a highly stringent mismatch detection mechanism. Although it might be argued that FnCas9 shows a generally lower efficiency of DNA cleavage due to which off-target cleavage events are diluted, the case of *c-MYC* where it exhibits comparable cleavage efficiency as SpCas9 yet maintaining no cleavage at off-target loci rules out such an explanation (Fig. 5B). Taken together, in our studies FnCas9 did not show any off-target activity, a prerogative for therapeutic gene targeting.

### FnCas9 mediated genome editing shows a higher efficiency of genetic insertions as compared to SpCas9

In a previous study (19), it was reported that FnCas9 produces a staggered pattern of DNA cleavage with a predominantly 4-nt 5’ overhang in the target strand. Interestingly, similar properties of staggered DNA cleavage have also been reported for Cpf1 protein which recognizes a T-rich PAM, generates sticky ends and is less tolerant to single or double mismatches in the crRNA sequence (33). In addition, Cpf1 has also been associated with very low off-targeting in mammalian cells (34). Observing certain parallels in the mode of cleavage between Cpf1 and FnCas9, we first examined the cleavage site of FnCas9 for multiple targets (*GFP, EMX1, VEGFA3* and *HBB*). We cloned sticky ended products of FnCas9 mediated *in vitro* cleavage reaction into a destination vector and performed bidirectional Sanger sequencing of the clones. We observed that in nearly all clones, the target strand was cleaved at 3bp upstream of the NGG PAM. However, the non-target strand showed different positions of cleavage (3-8 bp away from the PAM) depending upon the sequence being cleaved (Fig. 5D). In contrast, SpCas9 showed cleavage 3bp upstream of the PAM for the non-target strand as reported previously (8, SI Appendix, Fig. S4B). Taken together, these experiments reveal a flexible non-target strand cleavage activity by FnCas9 which has not been seen so far in other naturally occurring Cas9 proteins. Single molecule studies would be required to dissect how FnCas9 maintains this flexibility in non-target strand cleavage and the conformational changes that it encounters upon binding to a target.

We speculated that the sticky ends generated by FnCas9 might impact the rate of homology directed repair (HDR) mediated genomic insertions. Since HDR is an overarching component of therapeutic genome editing, we interrogated if FnCas9 can integrate foreign DNA in the form of a GFP reporter in a targeted fashion. We transfected HEK293T cells with a DNA template containing an out-of-frame GFP sequence flanked by homology sites to the mammalian *FASN* locus along with SpCas9 or FnCas9 with corresponding sgRNAs. A scrambled sgRNA sequence was used as control for both. We sorted cells on the basis of transient GFP expression from Cas9 containing plasmids after 72 h and allowed them to grow for 10 d to generate in-frame GFP from the endogenous FASN locus targeted by HDR. After 10 d, we analyzed the levels of the endogenous GFP by FACS. We consistently observed higher (~4.6 fold) GFP positive cells in FnCas9 transfected cells compared to SpCas9 transfected cells suggesting that FnCas9 produces higher efficiency of genetic insertions possibly through HDR (Fig. 5E). Using genotyping primers, we successfully validated these HDR events at the targeted locus (Fig. 5E). Collectively, these results suggest that FnCas9 shows higher levels of site-specific genomic insertions in HEK293T cells.

### FnCas9 generates detectable levels of genome editing in patient-derived iPSCs

Therapeutic genome editing trials for hemoglobinopathies have so far relied on ex-vivo gene editing strategies in patient derived hematopoietic or induced pluripotent stem cells. Intrigued by the specificity and higher HDR efficiency of FnCas9 mediated genome editing and to examine its efficacy in a clinical setup, we next tested its activity in a disease model of primary cells (Fig. 6A). To this end, we collected peripheral blood mononuclear cells from a patient carrying the homozygous sickle cell anemia mutation and generated induced pluripotent stem cells (iPSCs) using Sendai virus mediated reprogramming (SI Appendix, Fig. S4C). We characterized the resultant iPSC colonies with respect to their characteristic dome shaped morphology and expression of pluripotency markers OCT4 and SOX2 (Fig. 6B). To further establish their pluripotent character, we differentiated the iPSCs using defined factors into the three lineages and validated them by successful expression of ecto, meso and endodermal marker proteins SOX1, α-SMA and GATA4 respectively (SI Appendix, Fig. S4D). Digital karyotyping revealed no abnormal chromosomal aberrations in these iPSCs at different passages (SI Appendix, Fig. S4E).We then proceeded to electroporate these cells with FnCas9 or SpCas9. Since CRISPR clinical trials are currently focused on RNP mediated gene editing due to shorter persistence of the CRISPR complex and subsequent lesser off-targeting inside the cell (35), we generated RNP complexes targeting the mutant sickle cell anemia locus. For targeting the HBB locus harboring the sickle cell mutation, we attempted two different strategies: a single stranded donor nucleotide (ssODN) with 50bp homology-arms containing the WT beta globin gene or a double stranded DNA fragment carrying the same sequence. We electroporated patient iPSCs with RNP complexes carrying FnCas9/SpCas9-GFP along with donor templates, sorted GFP positive cells and analyzed the targeting locus using next generation sequencing. Consistent with reports in literature of very low targeting efficiency in iPSCs without the use of AAV donor vectors (36), we observed very few HDR events in both SpCas9 and FnCas9 over the scrambled control. Strikingly the number of significant HDR events was higher for FnCas9 as compared to SpCas9 when an ssODN was used (Fig. 6C). This is similar to what is reported for Cpf1 where the staggered cleavage on DNA has been attributed to a higher incorporation rate of ssODNs (37) and a lower rate of incorporation seen when blunt ended oligos are used. Indeed, we observed almost no HDR events with FnCas9 when a blunt double stranded oligo was used even though SpCas9 showed very low but significant detectable HDR events at the target (SI Appendix, Fig. S4F). Taken together, these results suggest that HDR events can be generated by FnCas9 based editing, particularly with the use of ssODNs and this can be utilized for correcting disease causing mutations in patient derived cells.

**Figure 6.**
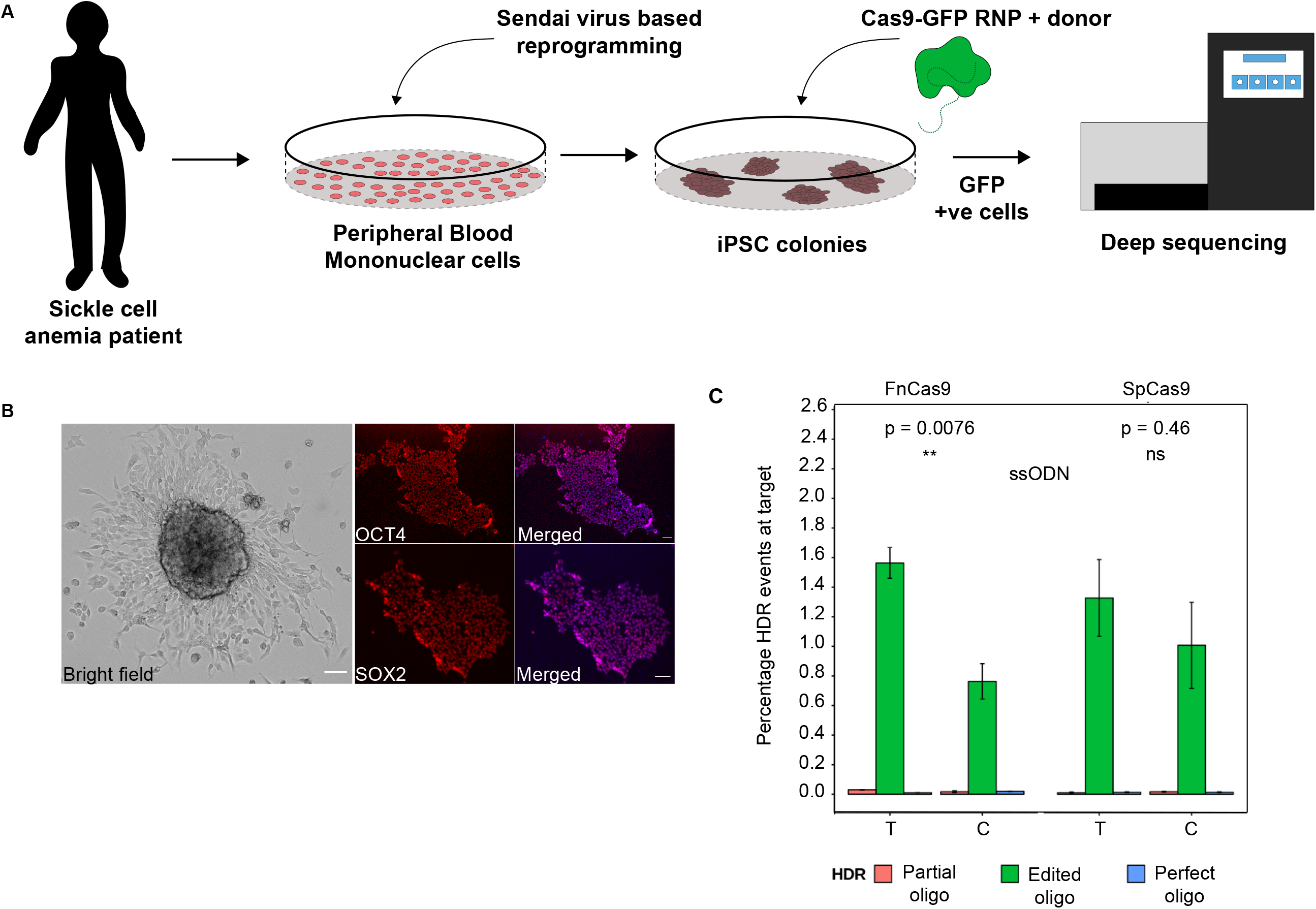
FnCas9 based genome editing in iPSCs. (A) Schematic showing genome editing using FnCas9 in patient derived iPSCs. (B) Micrograph showing an SCD patient derived iPSC colony (left). iPSC colony stained for pluripotency markers (Oct4 and Sox2) and Merged image with DAPI (in blue) is shown on the right. Scale bars 25 um. (C) HDR based correction of the sickle cell mutation in patient iPSCs using ssODN donor shown as percentage of HDR events at the target. Error bars represent SEM (3 independent experiments). Types of HDR detected are represented below. T = Test, C = Control.

## DISCUSSION

Precise genome engineering depends on the ability of the genome-editing agent to interrogate a given DNA loci with high specificity and minimum off targeting (38). The bacterial type II CRISPR Cas system has been efficiently exploited to edit a variety of genomes and has recently gained prominence for gene therapy applications, predominantly for monogenic disorders like beta thalassemia and sickle cell anemia (39–40). As one of the first Cas9 proteins described, SpCas9 has been highly successful in gene editing in diverse biological systems but reports of off-targeting remain a concern, particularly those of large deletions and complex rearrangements (41). In this study, we show that FnCas9, a Type II orthogonal Cas9 protein shows very high specificity to its target and negligible affinity with off-targets thereby increasing the repertoire of naturally occurring Cas9 proteins that can be used for therapy.

Binding studies of FnCas9 with its intended targets and off-targets reveal that FnCas9 has a fundamentally different mode of negotiating off-targets with PAM distal mismatches. Whereas highly specific engineered domains of SpCas9 remain in an associated ‘cleavage incompetent’ state with its off targets (24), FnCas9 shows greatly reduced affinity to such substrates possibly by eviction from the target upon encountering mismatches. Since the REC3 domain, which determines this specificity in the engineered variants is structurally unique in FnCas9, further biophysical studies aimed at visualizing the mode of action of this domain might elucidate the exact nature of target discrimination.

Systematic dissection of FnCas9 cleavage across different substrates reveal that its activity is largely abolished in the presence of 2 mismatches, most prominently at the PAM distal end. This understanding sets up a framework for FnCas9 sgRNA design algorithms with a high degree of specificity by rationally introducing mismatches in sgRNA sequences at defined positions that can distinguish between the target and a very similar off-target.

Although FnCas9 has a large size that can potentially deter its genomic interrogation to various sites, we observed successful *in vivo* genome editing at each of the genomic loci interrogated suggesting that it can access endogenous targets once it is inside the nucleus. Smaller sized Cas9 proteins offer the advantage of ease of delivery but can be more promiscuous in binding to unintended sites in the genome highlighting the importance of a balance between size and specificity.

In our cellular studies, FnCas9 showed negligible off-targeting at the various loci tested. Although certain engineered Cas9 proteins have shown very high discrimination of off-targets in the genome, to our knowledge this property is not seen among naturally occurring Cas9 proteins. How FnCas9 has evolved into a more precise genome editor than its other bacterial orthologs is a matter of further study. A study of structurally similar proteins from other species of bacteria and archaea might reveal genome editing proteins with beneficial attributes.

Finally, we show that FnCas9 can be used for HDR based modification of a desired target in the genome. The generation of staggered DNA cleavage, similar to Cpf1, is unique among naturally occurring Cas9 proteins and opens up the possibility of novel mechanisms of nuclease domain engagement with the target and non-target strands. Sticky ends might also explain higher rates of HDR seen in cells, a highly attractive feature of FnCas9 that can be suitably exploited in genetic engineering of a target loci and lays the foundation for including this protein to currently available genome editing agents for potential gene therapy applications. Although we did not achieve high levels of gene editing in patient-derived iPSCs using FnCas9 RNPs, delivery of both the protein and donor can be vastly improved using AAV vectors as reported in some recent studies (36). The bulky nature of the protein will necessitate shorter engineered versions that will be more apt for delivery, potentially by removal of redundant domains yet retaining the on-target specificity of FnCas9. Due to the modularity of the Cas9 enzyme, such constructs would need to exploit the steadily evolving functional link between REC3 and substrate mismatch discrimination.

Our studies indicate that FnCas9 might dissociate from its mismatched substrates rather rapidly resulting in minimal affinity with off-targets, a property that is strikingly dissimilar from other rationally designed Cas9 proteins that are highly specific (24). Remaining associated with off-target loci might result in transcriptional repression in these regions that can confound on-target activity and lead to cleavage-independent gene expression outcomes. Although it remains to be tested, FnCas9, due to its low binding affinity to off-targets, might present a more attractive scenario for genome editing where even long term persistence of the FnCas9 inside the cell might not induce off-targeting because of the high threshold for DNA binding when mismatches to the targets is encountered.

FnCas9, a type II-B Cas system has evolved separately from the II-A and II-C systems comprising of the majority of the commonly used Cas9 proteins used for genome editing (42). Its distinct structural attributes leading to high targeting specificity hints at the existence of more members of the II-B family that can be similarly explored for mammalian genome editing. Further, FnCas9 can be structurally engineered to render it competent for base editing, an attractive therapeutic application for monogenic disorders where the current generation of base editors have so far shown variability in off-targeting (43).

## MATERIALS AND METHODS

### Plasmid Construction

The gene encoding full length *Francisella novicida* (Fn) Cas9 nuclease residues 1-1629 bp) was PCR amplified using PX408 (Addgene 68705) as a template and cloned in pET28-His-10-Smt3 vector (a kind gift from Prof. Stewart Shuman and Dr. K.M. Sinha) and pET-His6-GFP-TEV-LIC vector (Addgene 29663) following restriction enzyme based cloning and ligation independent cloning (LIC) respectively. Catalytically inactive FnCas9 double mutants were generated on pET-His6-FnGFP-TEV-LIC (this paper) plasmid backbone by QuickChange II site directed mutagenesis kit (Agilent) following manufacturer’s protocol with some modifications.

For the expression of SpCas9 and FnCas9 in mammalian cells, 3xHA-SpCas9, 3xHA-FnCas9 and FnCas9 BbsI sgRNA cloning site were synthesized as gene blocks (GenScript) and cloned into PX458 (Addgene 48138) backbone by restriction enzyme-based cloning. FnCas9 BbsI sgRNA cloning site was cloned in PciI and XbaI sites of PX458 to make it suitable for FnCas9 sgRNA expression and in this vector (PX458-3xHA-FnCas9) 3xHA-FnCas9 gene block was cloned in AgeI and FseI sites. Desired guide RNA sequences were cloned as annealed oligonucleotides with appropriate overhangs in BbsI sites following the established protocol (44). A complete list of sgRNA target sites is available in SI Appendix, Dataset S1. All the constructs were sequenced before being used.

### Protein purification

The proteins used in this study were purified as reported previously (8). Briefly, plasmids for different Cas9 proteins were expressed in *Escherichia coli* Rosetta2 (DE3) (Novagen). The protein expressing Rosetta2 (DE3) cells were cultured at 37°C in LB medium (supplemented with 50mg/l Kanamycin) until OD_600_ reached 0.6 and protein expression was induced by addition of 0.5mM isopropyl β-D-thiogalactopyranoside (IPTG). The Rosetta2 (DE3) cells were further cultured at 18°C overnight and harvested by centrifugation. The *E.coli* cells were resuspended in lysis buffer (20mM HEPES, pH 7.5, 500mM NaCl, 5% glycerol supplemented with 1X PIC (Roche), 100ug/ml lysozyme and lysed by sonication and centrifuged. The lysate was affinity purified by Ni-NTA beads (Roche) and the eluted protein was further purified by size-exclusion chromatography on HiLoad Superdex 200 16/60 column (GE Healthcare) in 20 mM HEPES pH 7.5, 150 mM KCl, 10% glycerol, 1mM DTT. The concentration of purified proteins were measured by Pierce BCA protein assay kit (Thermo Fisher Scientific).The purified proteins were stored at −80°C until further use.

In case of 6X His-MBP-dSpCas9, the 6X His-MBP tag was removed by incubating the affinity bound protein with PreScission Protease overnight in cleavage buffer (50mM Tris-Cl, pH8, 150mM NaCl, 1mM EDTA,1mM DTT). The cleaved Cas9 protein was separated from fusion tag on HiLoad Superdex 200 16/60 column (GE Healthcare) in 20 mM HEPES pH 7.5, 150 mM KCl, 10% glycerol, 1mM DTT.

### Microscale thermophoresis (MST) assay

MST assay was performed following earlier mentioned protocol (45–46). dSpCas9-GFP and dFnCas9-GFP proteins were used along with PAGE purified respective IVT sgRNAs. Note that IVT sgRNAs were purified by 12% Urea-PAGE. The binding affinities of the Cas9 proteins and sgRNA (ribonucleoprotein) complexes towards different genomic loci like c-Myc promoter, EMX1, VEGFA3 were calculated using Monolith NT. 115 (NanoTemper Technologies GmbH, Munich, Germany). RNP complex (Protein:sgRNA molar ratio, 1:1) was reconstituted at 25°C for 10 mins in reaction buffer (20 mM HEPES, pH7.5, 150mM KCl, 1mM DTT, 10mM MgCl_2_). HPLC purified 30 bp dsDNA (IDT) of different genomic loci with varying concentrations (ranging from 0.09nM to 30μM) were incubated with RNP complex at 37^0^ C for 30 min in reaction buffer. The sample was loaded into NanoTemper standard treated capillaries. Measurements were performed at 25°C using 20% LED power and 40% MST power. All experiments were repeated at least two times for each measurement. All Data analyses were done using NanoTemper analysis software.

### Electrophoretic mobility shift assay (EMSA)

EMSA was performed following the protocol reported earlier (47). 250nM sgRNA was incubated with different concentrations of catalytically inactive dead (D11A-H96A FnCas9) Cas9 at 25°C for 10 min in reaction buffer (20 mM HEPES, pH7.5, 150mM KCl, 1mM DTT, 10% glycerol, 10mM MgCl_2_). The samples were resolved by 8% Native polyacrylamide gel electrophoresis (0.5X TBE buffer with 2mM MgCl_2_) run at 4°C and the gel was stained by SYBR Gold Nucleic acid Gel stain (ThermoFisher Scientific) for 30 min at room temperature in shaking condition. RNA was visualized by Typhoon FLA 7000 (GE Healthcare) and quantified by ImageJ.

### CASFISH

CASFISH was done following standard CASFISH protocol (48). Desired number of cells were seeded on 22×22 mm 0.1% Gelatin (Sigma)-coated coverslips and 24 hrs post-seeding, cells were fixed at −20°C for 20 min in a pre-chilled solution of methanol and acetic acid at a 1:1 ratio. Fixed cells were washed thrice for 5 min each with 1X PBS with gentle shaking followed by incubation for 30 min at 37°C in blocking buffer (20mM HEPES pH 7.5, 150mM KCl, 10mM MgCl_2_, freshly added 1mM DTT, 5% (vol/vol) Glycerol, 2% BSA, 0.1% Triton X). CASFISH probes were assembled by mixing 200mM dCas9-GFP with 200mM sgRNA targeting Telomeric region (1:1 molar ratio) in blocking buffer and incubated at room temperature for 10 min. The assembled RNP complex was applied on pre-blocked cells and incubated for 2-4 hrs in humid chamber kept at 37°C. The reaction was terminated by removing RNP complex solution followed by washing thrice with blocking buffer. Washed samples were stained with 5μg/ml DAPI and were mounted on glass slides before imaging. CASHFISH samples were imaged on a DeltaVision Ultra microscope (GE Healthcare, software Acquire Ultra 1.1.1) equipped with 60X oil-immersion objective (NA 1.4). Images were processed using ImageJ.

### Comparative structural study between Fn- and Sp- cas9

#### Electrostatic analysis

For the purpose of understanding the underlying electrostatic contributions in the Fn- and Sp- cas9 structures, we used the standalone version of PDB2PQR (49–50) to assign charge and radius parameters for CHARMM force field to the Fn- (PDBID: 5B2O) and Sp- (PDBID: 5F9R) crystal structures.

The electrostatics was calculated using Adaptive Poisson-Boltzmann Solver (APBS) (51). APBS uses iterative solvers to solve the nonlinear algebraic equations resulting from the discretized Poisson-Boltzmann equation with a fixed error tolerance of 10^-6. The values are coloured in the range of −10 to 35 k_b_ T e_c_^−1^ where

- k_b_ is Boltzmann’s constant: 1.3806504 × 10^−23^ J K^−1^
- T is the temperature of your calculation in K
- e_c_ is the charge of an electron: 1.60217646 × 10^−19^ C

The images were generated using UCSF Chimera tool.

#### Contact analysis

To study the interaction between the nucleotide and protein, atomic contacts were calculated using a 4 Å cut-off. The graph was generated using MATLAB.

### Derivation of PBMCs from Sickle Cell Anemia Patient

The present study was approved by Ethics Committee (Ref no. CSIR/IGIB/IHEC/17-18) and Institutional Committee for Stem Cell Research (Ref no. IGIB/IC-SCR/9), Institute of Genomics and Integrative Biology, New Delhi, India. Written informed consent was obtained from the sickle cell patient and blood was collected. Peripheral blood was collected in heparin coated vacutainers from sickle cell anemia patient (6 yrs, Male). Mononuclear cells were isolated from 6ml blood sample using standard Ficoll-Paque gradient centrifugation (52). Briefly, heparinized blood was diluted 1:1 in cell culture grade Dulbecco’s PBS (DPBS) (Gibco) and was layered over Ficoll-Paque PLUS (density 1.077 g/mL, GE Healthcare) in 15ml falcon tube. The tubes were centrifuged for 330 x g for 30 minutes at room temperature with brake-off. The cell interface layer (buffy coat) was harvested and washed twice with 10% FBS (ES qualified) in DPBS solution for 15 min at 330 xg. PBMCs were resuspended in StemPro™-34 SFM Medium supplemented with L-Glutamine to a final concentration of 2 mM and cytokines with recommended final concentrations, SCF (C-Kit Ligand) Human Recombinant Protein (Cat. No. PHC2111) 100 ng/mL; FLT-3 Human Recombinant Protein (Cat. No. PHC9414) 100 ng/mL; IL-3 Human Recombinant Protein (Cat. No. PHC 0034) 20 ng/mL and IL-6 Human Recombinant (Cat. No. PHC0034) 20 ng/mL. 5 x 10^5^ cells/ml were plated in four wells of a 24 well plate and maintained in standard culture conditions (37°C, 5% CO2) and feeded daily for next 3 days.

### Generation of iPSCs and maintenance

Four days after initial culture, PBMCs were reprogrammed using CytoTune™-iPS 2.0 Sendai Reprogramming Kit (Thermo Fisher Scientific) containing three SeV vectors encoding *OCT3/4, SOX2, KLF4*, and *c-MYC* (53). Briefly, volume of each viral vector for the transduction mix was calculated as per manufacturer’s protocol and added to 3 x 10^5^ cells in 1ml PBMC medium followed by centrifugation at 1000 x g for 30 minutes at room temperature. Cells were resuspended and plated in 2ml media in 1 well of a 12-well plate. 2 days post-transduction, cells were seeded on Matrigel (Corning) coated 6-well culture plates in 2ml StemPro™-34 SFM Medium without cytokines (54–55). Further after 4 days, cells were transitioned to mTeSR™1 (STEMCELL Technologies) medium. After 2 weeks, iPS colonies were picked and transferred to Matrigel coated 12-well plates. SCD-patient derived iPS cell line was maintained in mTeSR™1 medium and subcultured using enzyme free passaging reagent, ReLeSR™ (STEMCELL Technologies). Mutation in patient derived-iPSCs were confirmed using Sanger sequencing.

### Cas9 Genome Editing in iPSCs

iPSCs were treated with 10 μM ROCK inhibitor (Y-27632) 1 hour before electroporation. 70%-80% confluent iPSC colonies were harvested using StemPro™ Accutase™ Cell Dissociation Reagent (Gibco) and pipetted to make single cell suspension. 5ul volume of RNP complex mix was made in 1:3 molar ratio in Resuspension Buffer R for each electroporation, and incubated for 20 minutes at RT. 3.5 x 10^5^ cells were resuspended in 10 μL of Resuspension Buffer R per electroporation condition and 20μM ssODN or 1ug dsDNA with 5ul of complete RNP complex was added and electroporation performed using Neon^®^ Transfection System 10 μL Kit (Thermo Fisher Scientific) with a single pulse at 1200 V, 30 milliseconds pulse width (56). The electroporated cells were transferred immediately to a Matrigel coated 24 well plate containing 0.5 ml of mTeSR™1 with 10uM ROCK Inhibitor and incubated at 37°C humidified incubator supplied with 5% CO2.

After 12 hours, cells were washed and re-incubated with fresh mTeSR™1 medium with 10uM ROCK Inhibitor for 1 hr and then harvested. 25,000 GFP positive cells per sample were sorted using BD FACS Melody Cell Sorter (BD Biosciences-US) and re-plated. gDNA was isolated after 7 days using Wizard^®^ Genomic DNA Purification Kit (Promega) for preparation of 16S Metagenomic Sequencing Library.

### iPSC library preparation and sequencing

The 16S Metagenomic sequencing library preparation protocol was adapted for library preparation. Briefly, HBB locus was amplified using forward and reverse primers along with overhang adapter sequences using Phusion High-Fidelity DNA polymerase (Thermo Fisher). AMPure XP beads (A63881, Beckman Coulter) were used to separate out amplicons from free primers and primer dimmers. Dual indexing was done using Nextera XT V2 index kit followed by another round of bead based purification. The libraries were quantified using Qubit dsDNA HS Assay kit (Invitrogen, Q32853) and were also loaded on agarose gel for qualitative check. Libraries were normalized, pooled and was loaded onto illumina MiniSeq platform for a 150bp paired end sequencing run.

### Data Analysis

Determination of indel frequency from sequencing data was performed using CRISPResso2 v2.0.29 (57) with the following parameters ‘--ignore_substitution -- min_paired_end_reads_overlap 10 --max_paired_end_reads_overlap of 500. We have detected by sanger sequencing that the cleavage positions for FnCas9 on the non-target strand varied from 3-8bp upstream of the PAM, so, quantification window for indel detection was set 3 to 8 bp upstream of PAM. For c-Myc samples (ontarget and OT1) amplicon sequences were shorter (165 and 171 respectively) than the sequencing read length of 2X250bp so, 50 bp were cropped from the end of reads to achieve the required minimum homology (60% according to CRISPResso2) with the amplicon. CRISPR-DAV has been used with default parameters (58) for HDR analysis of sickle cell IPSCs.

## ACKNOWLEDGMENTS

This work was supported by Council for Scientific and Industrial Research (CSIR) Mission Mode Project grant HCP0008 to D.C. and S.M. The *MYC* construct was a kind gift from the lab of Shantanu Chaudhury, CSIR IGIB and the FASN construct used for HDR experiments was generously shared by Mihail Sarov, MPICBG, Dresden. The authors wish to thank the lab of Frank Buchholz and Beena Pillai for constructs and cell lines used in the study and Debasis Dash, CSIR IGIB, Sambit Dalui, CSIR IICB, and all members of S.M. and D.C. labs for helpful discussions and feedback. The support of Manish Kumar, Imaging facility CSIR IGIB, Manish Kumar Rai and Nanotemper, Bangalore (MST experiments) is also gratefully acknowledged.

## AUTHOR CONTRIBUTIONS

D.C., S.M., S.A., A.M., D.P., and M.Az. designed the experiments. S.A., A.M., D.P., M.Az, N.S., M.Ai., D.S., S.S. and S.J. performed experiments and collected data, A.H.A., D.P., A.M. and A.R. analyzed data, M.K., R.R., S.J. and S.R. helped with developing iPSC lines, D.C. wrote the manuscript. All authors reviewed the manuscript and approved the conclusions.

## DECLARATION OF INTERESTS

The authors declare no conflict of interest.

## Notes

#### Summary of Updates

This version of the manuscript has been revised to include more data on invivo genome editing and also includes additional authors. The format of old figures have been changed and some data has been updated in the light of new results.

