## Supplemental Information for "*Francisella novicida* Cas9 interrogates genomic DNA with very high specificity and can be used for mammalian genome editing"

**This PDF file includes:**

Supplementary text
Figs. S1 to S4

### Supplementary Information Text

#### Methods

##### *In vitro* transcription:

*in vitro* transcribed sgRNAs were synthesized using MegaScript T7 Transcription kit (Thermo Fisher Scientific) using T7 promoter containing template as substrates. IVT reactions were incubated overnight at 37°C followed by NucAway spin column (Thermo Fisher Scientific) purification. IVT sgRNAs were stored at -20°C until further use.

##### *In vitro* Cleavage assay (IVC):

Cas9-sgRNA complex was reconstituted at 25°C for 10 mins in reaction buffer (20 mM HEPES, pH7.5, 150mM KCl, 1mM DTT, 10% glycerol, 10mM $MgCl_2$). The required amount of desired DNA substrates (given in the respective figure legends) were incubated in reaction buffer with reconstituted RNP complex at 37°C. The details of different experiments were given in respective figure legends. The samples were treated with 1ul of 20mg/ml Proteinase K in 15 ul reaction for 15 mins at 55°C. The reaction products were resolved in EtBr-stained 1% agarose gel, visualized by Syngene UV transilluminator and quantified by ImageJ. Note that respective substrates used for IVC assay were mentioned in corresponding figure legend and the reactions were stopped by adding 25 mM EDTA final concentration before addition of Proteinase K in kinetics experiment.

##### Cloning of IVC products:

500nM of Cas9-sgRNA complex was reconstituted at 25°C for 10 mins in reaction buffer with 500ng of PCR product for each locus. *In vitro* cleaved products were gel eluted using QIAquick Gel extraction kit (Qiagen). Gel purified products were then cloned in pJET 1.2/blunt cloning vector using sticky-End cloning protocol of

CloneJET PCR Cloning kit (Thermo Fisher Scientific). Recombinant pJET 1.2/blunt cloning vector was transformed into chemically competent DH5α *E. coli* cells. Transformed colonies were inoculated in LB media containing 100ug/ml ampicillin and purified using QIAprep spin Miniprep kit (Qiagen). Purified plasmids were screened by PCR and Sanger sequencing was performed (AgriGenome Labs). For FnCas9, 37 clones for EMX1, 41 for VEGFA3, 46 for GFP and 40 for HBB were sequenced. As a control 38 clones of HBB cleaved by SpCas9 were also sequenced.

**Cell culture and transfection:**

HEK 293T, Neuro-2A and Neuro-2A-GFP cells were grown in DMEM media supplemented with high glucose (Invitrogen), 2mM GlutaMax, 10% FBS (Invitrogen), 1X antibiotic and antimycotic (Invitrogen) at 37°C in 5% $CO_2$. Mouse embryonic stem cells (R1/E) were cultured in DMEM media (Invitrogen) supplemented with 20% Pansera, 1% PenStrep (Invitrogen), 0.1% B-ME, 1% NEAA (Invitrogen) and 40ug/mL LIF (MPI CBG). Transfections of mammalian cells were performed using Lipofectamine 3000 Reagent (Invitrogen) following manufacturer's protocol.

**ChIP assay:**

Neuro-2A-GFP cells on 10cm dishes were transfected with 50μg of FnCas9 (Addgene #68705) plasmid. 24hrs post transfection, 50μg of respective IVT sgRNAs were transfected. Cells were harvested 24 hours post sgRNA transfection. Target cells for ChIP assays were cross-linked with 1% formaldehyde (Sigma) followed by vortexing and incubation with rotation at room temperature for 15 mins and subsequently quenched by addition of 125mM glycine. Supernatant was discarded and the pellet was dissolved in 5ml Lysis Buffer I (50mM HEPES-KOH pH 7.5, 140mM NaCl, 1mM EDTA, 10% Glycerol, 0.5% NP-40, 0.25%TritonX-100, 1X Roche complete Protease inhibitor) followed by centrifugation at 1350xg for 5 mins at 4°C. After discarding supernatant, pellet was resuspended in 5ml Lysis Buffer II (10mM Tris-Cl pH 8, 200mM NaCl, 1mM EDTA, 1X Roche protease inhibitor), rotated at 4°C for 10mins, followed by centrifugation at 1350xg for 5 mins at 4°C. The nuclear pellet was resuspended in sonication buffer (20 mM Tris-Cl pH8, 150 mM NaCl, 2mM EDTA, 0.1% SDS, 1% TritonX-100, 1xRoche protease inhibitor) for sonication (3xHigh + 2xLow, each 30secs OFF and 30secs ON cycle). 50ul (30ug/ul conc) of Dynabeads Protein G per reaction was taken. Beads were washed twice with Ab binding and washing buffer and then incubated with Salmon sperm DNA (AM 9680) for 2 hrs. The cell lysate was allowed to bind with 10ug of anti-HA Ab (Abcam ab9110) for O/N at 4°C. 1% of the total volume was kept as Input. Dynabeads were allowed to bind to the sample (incubated with Ab O/N) for 4-5 hrs at 4°C. Beads were washed the next day followed by sample elution and reverse crosslinking was done for both input and IP samples keeping them at 65°C O/N. RNase treatment (0.2mg/ml) was done at 37°C for 2hrs, followed by ProteinaseK (0.2mg/ml) treatment at 55°C for 45 mins. DNA purification was performed with QIAquick PCR Purification Kit (Qiagen) and proceeded for qRT-PCR.

**Amplicon sequencing:**

HEK293T cells on six well dishes were transfected with 5µg of SpCas9 and FnCas9 containing sgRNAs targeting c-Myc, EMX1 and HBB. Cas9 plasmids with scrambled sgRNA were used as controls. 48 hours post-transfection GFP-positive cells were sorted by BD FACS Melody (BD Biosciences). Wizard Genomic DNA purification kit (Promega Corporation) was used to isolate genomic DNA. PCR primers were designed flanking the predicted double stranded break site using Phusion High-Fidelity DNA polymerase (Thermo Fisher Scientific). The DNA samples were quantified using Qubit dsDNA HS Assay kit (Invitrogen). Library preparation was done using NEB Next Ultra DNA Library Prep Kit for Illumina (NEB). First, the amplicons were end repaired and mono-adenylated at the 3' end. Next, NEB hairpin-loop adapters were ligated to the DNA fragments in a T4-DNA ligase-based reaction. Following ligation, the loop containing Uracil was linearized using USER Enzyme (a combination of UDG and Endo VIII) to make it available as a substrate for PCR based indexing in the next step. Barcodes incorporated during PCR using unique primers enabled multiplexing. Library preparation followed by 250 bp paired end sequencing was done on illumina MiSeq platform by MedGenome Labs Pvt. Ltd. (Bangalore, India).

**Homology Directed Repair (HDR):**

1ug of sgRNA specific to FASN loci cloned in the same vector as Cas9 and 1 ug of GFP containing donor plasmid were electroporated in $1X10^5$ HEK293T cells in a 24-well format using 10ul tip of Neon Transfection System (ThermoFisher Scientific). Conditions for electroporation was 950V, 30 ms and two pulses. Cells were sorted 72 hours post-electroporation using BD FACS Melody (BD Biosciences) and sorted cells were allowed to grow for 10 days. Second FACS sorting was done to calculate the percentage of HDR efficiency against the respective Cas9 plasmids containing scrambled sgRNA and GFP donor plasmid. Sorted cells were harvested for genomic DNA isolation using Wizard Genomic DNA purification kit (Promega Corporation) from GFP sorted (+HDR) and non-GFP (-HDR) cells. PCR was performed with Herculase II Fusion DNA polymerase (Agilent) using GFP specific forward primer and FASN specific reverse primer located outside the right homology arm.

**Immunofluorescence staining of iPS cells to determine pluripotency:**

For live staining of pluripotent cells, TRA-1-60 Mouse anti-human, Alexa Fluor™ 594 Conjugate (Invitrogen) was used as per manufacturer's protocol. Briefly, 1:100 volume of the dye-conjugated antibody was directly added and mixed with cell culture medium and cells were incubated for 30 minutes at 37°C. Medium was aspirated and the cells were washed three times with FluoroBrite™ DMEM (Gibco) and imaging was done in this medium within 30 minutes.

For immunocytochemistry, cells were fixed in 4% paraformaldehyde for 15 min at room temperature. Fixed samples were washed three times with PBS and permeabilized with 0.25% Triton X-100 in PBS for 30 min at room temperature. Cells were blocked with blocking buffer (0.3% Triton-X, 10% Goat Serum and 1% BSA in PBS), washed three times with wash buffer (0.1% BSA and 0.1% Tween in PBS) and then incubated with primary antibodies diluted in 1% BSA, 1% Goat Serum and 0.25% Triton-X in PBS, overnight at 4°C. Primary antibodies were then washed three times with PBS and cells were incubated with secondary antibodies at room temperature for 1 hour. Cells were washed again and incubated with DAPI (Sigma) for 5 min at RT and further washed and stored in PBS at 4 °C until visualization. Fluorescence images were acquired using the EVOS FL Auto Cell Imaging System (ThermoFisher Scientific, USA).The list of the primary and secondary antibodies and dilution used is mentioned in SI Appendix, Dataset S1.

**Karyotype analysis:**
qPCR based genetic screening of SCD patient derived iPS cells was performed using hPSC Genetic Analysis Kit (STEMCELL Technologies) following the manufacturer's instructions. Results were analyzed using the application available at www.stemcell.com/geneticanalysisapp.

**Trilineage Differentiation of iPSCs:**

STEMdiff™ Trilineage Differentiation Kit (STEMCELL Technologies) was used for directed differentiation of iPSCs to the three germ layers. Briefly, for ectoderm lineage, 4 x $10^5$ cells were seeded in 0.5ml of STEMdiff™ Trilineage Ectoderm Medium with 10µM of Y-27632. For mesoderm and endoderm lineage, 5 x $10^4$ cells and 2 x $10^5$ cells were seeded in 0.5 ml of mTeSR™1 with 10µM Y-27632, respectively. Media was changed every other day with 1ml of appropriate STEMdiff™ Trilineage medium. Differentiating monolayer cultures with mesoderm and endoderm lineages were fixed 5 days after seeding and ectoderm lineage was fixed after 7 days. Lineage specific markers were analysed by immunocytochemistry.

**Quantification and statistical analysis:**

Gel image quantification was done using ImageJ. Statistical analysis and visualization has been done in R.

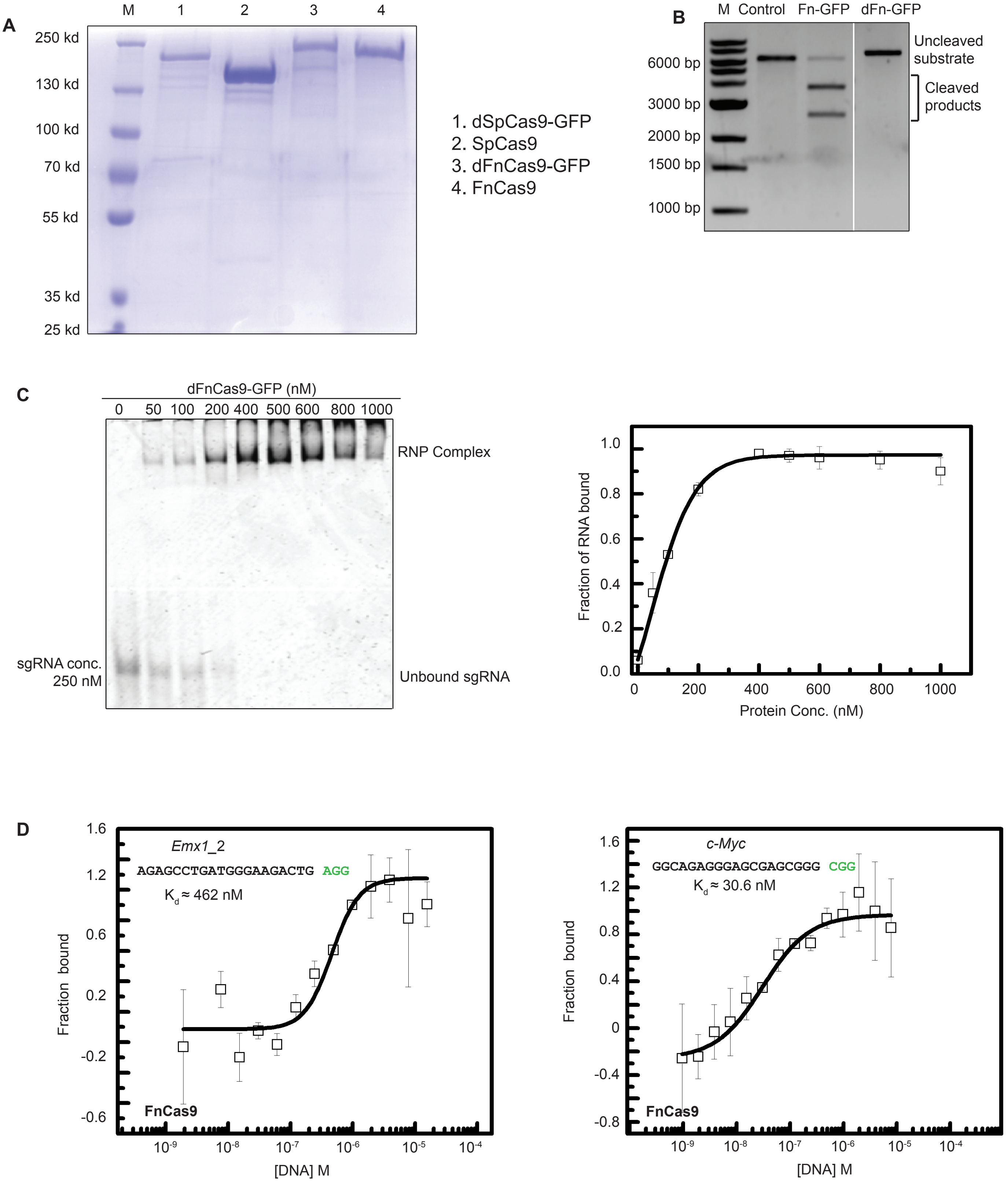


**Figure S1. FnCas9 shows different binding affinities with different substrates**

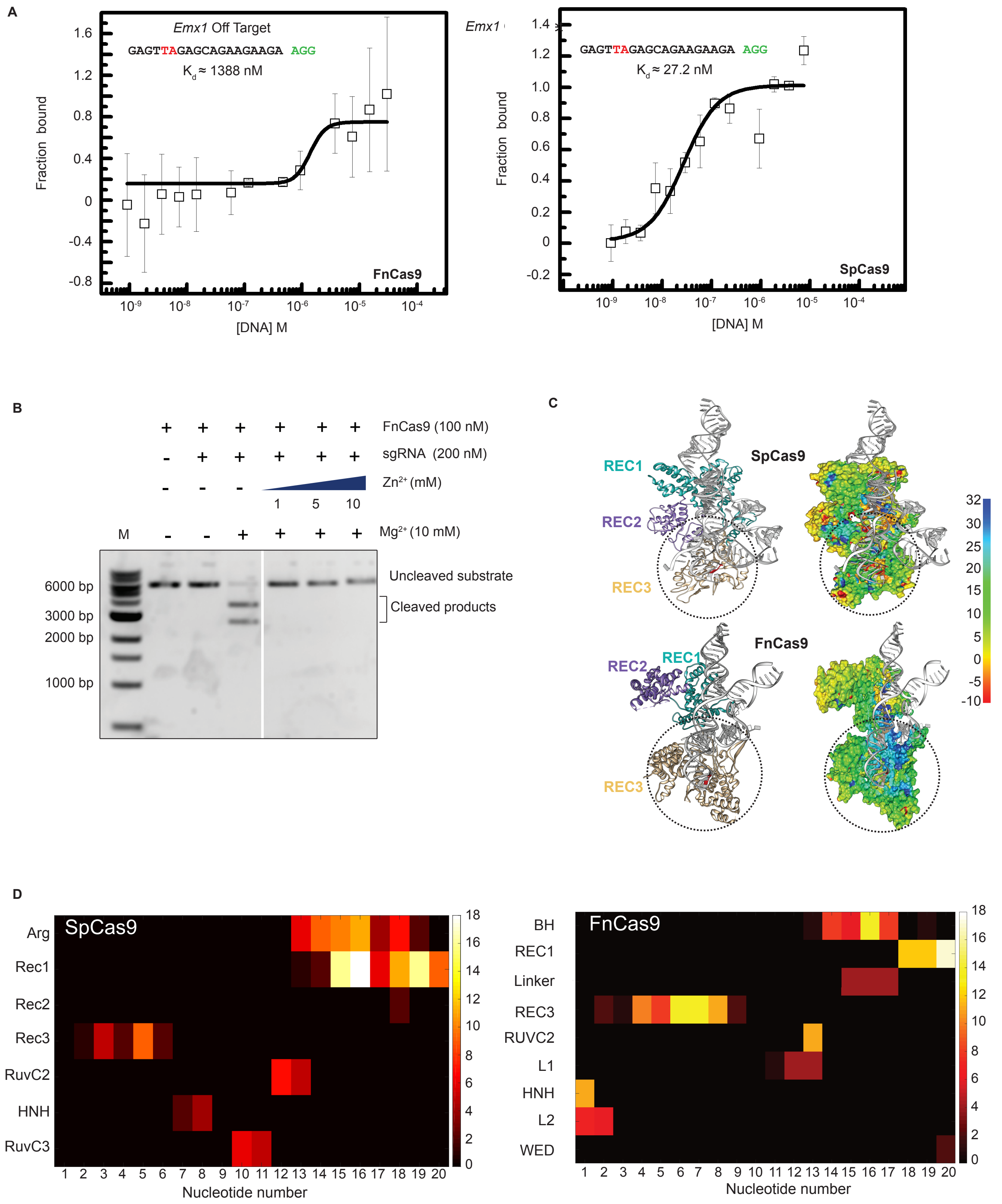


**Fig. S2. FnCas9 shows unique structural attributes for substrate binding and cleavage**

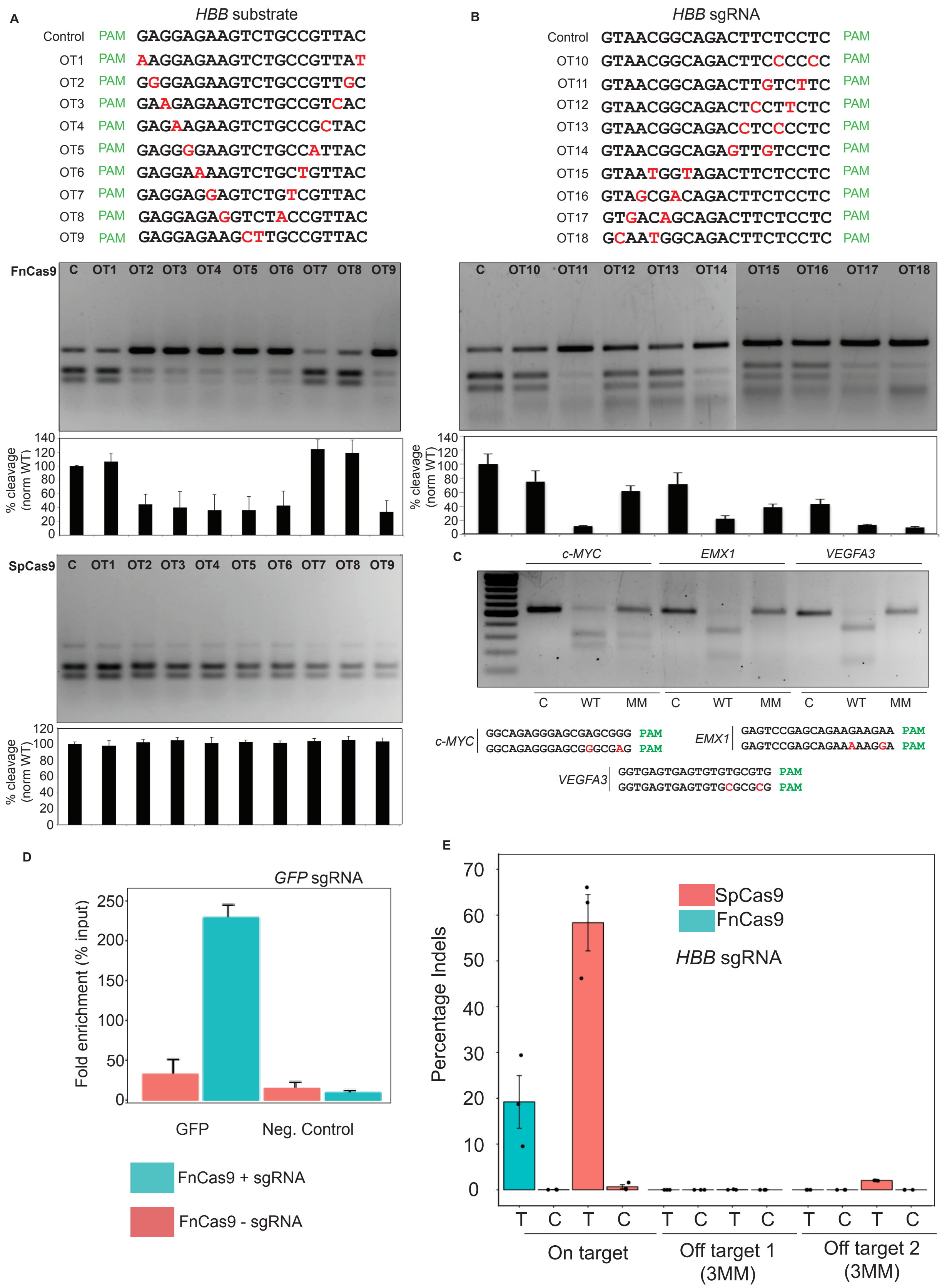


**Fig. S3. Cleavage activity of FnCas9 is affected in the presence of 2 mismatches in the substrate or the sgRNA**

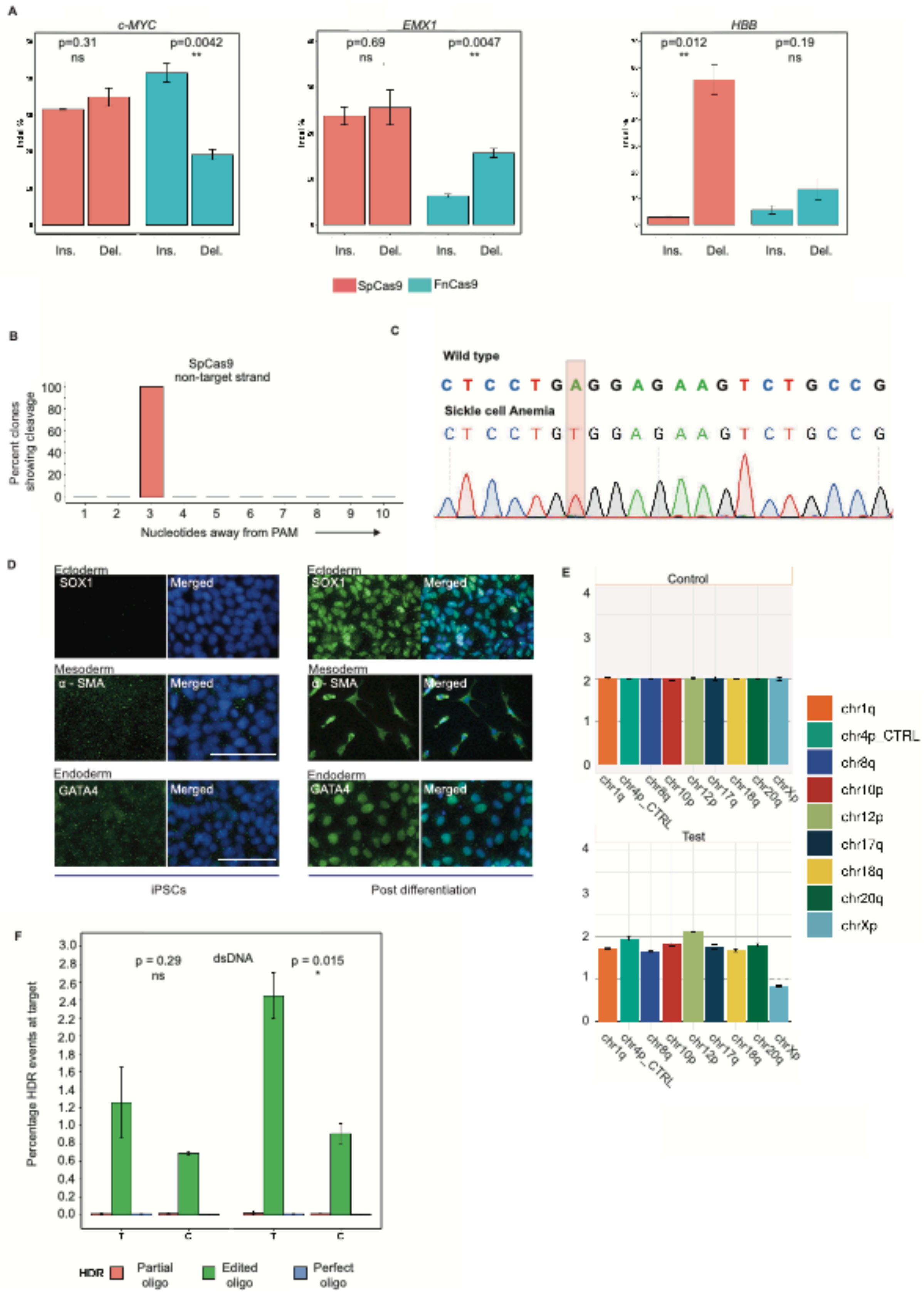


**Fig. S4. FnCas9 mediates *in vivo* genome editing in mammalian cells**

**Dataset S1**
Sheet 1 contains list of primers, oligos and sgRNAs used in the study, sheet 2 contains Antibodies used in immunostaining.

**Figure. S1. FnCas9 shows different binding affinities with different substrates**

(A) Coomassie gel showing different purified proteins used in the study.
(B) *In vitro* cleavage assay showing mutation of nuclease domains in FnCas9 abolishes its cleavage activity on a linearized plasmid (100 ng substrate, 50nM Cas9:150nM sgRNA).
(C) Left, representative EMSA showing binding of purified dFnCas9-GFP at different concentrations to its sgRNA sequence (250 nM). Right, quantification of protein: sgRNA binding. Error bars represent SEM (3 independent experiments).
(D) MST results showing binding outcomes of dFnCas9-GFP to *EMX1*_2 and *c-MYC* substrates expressed as normalized fluorescence units (y-axis) with respect to varying concentrations of purified substrate (x-axis). The substrate sequences are indicated in the box with the PAM shown in Green. Error bars represent SEM (2 independent experiments).

**Figure S2. FnCas9 shows unique structural attributes for substrate binding and cleavage**

(A) *In vitro* cleavage assays showing outcome of *c-MYC* substrate cleavage (2.5nM) with varying FnCas9 RNP concentrations (as indicated) across different time points. Note that cleavage activity increases with increasing concentration of FnCas9 RNP complex.
(B) MST results showing binding outcomes of dFnCas9-GFP (left) and dSpCas9-GFP (right) to *EMX1* off-target substrates expressed as

normalized fluorescence units (y-axis) with respect to varying concentrations of purified substrate (x-axis). The substrate sequences are indicated in the box with the PAM shown in Green. Red denotes the mismatches in the target. Error bars represent SEM (2 independent experiments).

(C) Activity of FnCas9 is abolished in the presence of $Zn^{2+}$ ions. *In vitro* cleavage assay showing FnCas9 is active on a substrate (linearized plasmid pCAGSS_GFP_RFP, 100ng) only in the presence of $Mg^{2+}$ and absence of $Zn^{2+}$.

(D) Crystal structures of FnCas9 (PDB 5B2O, top) and SpCas9 (PDB 5F9R, bottom) RNP in complex with substrate DNA in ribbon (left) and electrostatic potential surface representation (right). PAM is indicated in red and individual domains are color-coded. Heat-map of electrostatic contribution is indicated on the right. Note the higher positive electrostatic potential of FnCas9 domains (indicated by dotted circles) interacting with substrate. REC3 domain in each enzyme has been denoted with dotted circles.

(E) Quantification of electrostatic potential contribution of FnCas9 (top) and SpCas9 (bottom) on substrate DNA nucleotides. Nucleotide positions are indicated below from PAM proximal to PAM distal end (left to right).

**Figure S3. Cleavage activity of FnCas9 is affected in the presence of 2 mismatches in the substrate or the sgRNA**

(A) 2 Mismatches at different positions along the substrate alter FnCas9 cleavage outcomes while retaining SpCas9 activity. Representative *in vitro* cleavage outcome of HBB substrate (100ng) sequence showing mismatches at different positions (indicated in red) interrogated with FnCas9 (top) or SpCas9 (bottom) is shown (250nM RNP), PAM is indicated in green. Top band represent uncleaved target while the two bottom bands represent cleaved products. Quantification for each reaction is shown below the gel images. Error bars represent SD (3 independent experiments).

(B) Representative *in vitro* cleavage outcome of HBB sgRNA sequence showing mismatches at different positions (indicated in red) interrogated with FnCas9 is shown (substrate 100ng, RNP 250nM), position of PAM with respect to the sequence orientation is indicated in green. Top band represent uncleaved target while the two bottom bands represent cleaved products. Quantification for each reaction is shown below the gel image. Error bars represent SD (3 independent experiments).

(C) *In vitro* cleavage outcomes of *c-MYC*, *EMX1* and *VEGFA3* targets interrogated by WT FnCas9 showing abrogation of cleavage in the presence of 2 mismatches (substrate 100ng, RNP 250nM). PAM is represented in green and mismatched bases in red.

(D) Chromatin Immunoprecipitation (ChIP) of FnCas9 showing binding of FnCas9 RNP complex to endogenous GFP target in mouse Neuro-2A cell line. Fold enrichment (percentage of input) is shown on y-axis. A non-targeting region is taken as negative control. Error bars represent SEM (3 PCR replicates), data representative of two independent experiments.

(E) Indel events (expressed as percentage) obtained by amplicon sequencing upon SpCas9 or FnCas9 targeting *HBB* in HEK293T cells. Individual data points from independent experiments are shown. Error bars represent SEM (3 independent experiments). T = Test, C = Control.

**Figure S4. FnCas9 mediates *in vivo* genome editing in mammalian cells**

(A) Outcomes of SpCas9 or FnCas9 mediated genomic cleavage showing insertions/deletions (indels) per event at *EMX1*, c-*MYC* (right) and *HBB* loci detected after amplicon sequencing in HEK293T cells. Error bars represent SEM (3 independent experiments).

(B) Cleavage positions on non-target strand by SpCas9 determined using Sanger sequencing for *HBB* target. Y axis represents percentage of sequenced clones showing cleavage at a given nucleotide. X axis represents the position of the base away from PAM in the sgRNA.

(C) Chromatogram highlighting SNP in patient compared to wildtype (GAG>GTG).

(D) Representative micrographs showing expression of linage markers for the three germ layers, Ectoderm (SOX1), Endoderm (GATA4) and Mesoderm (α-SMA) in differentiated and undifferentiated states. DNA is stained with DAPI and represented in blue in merged images. Scale bar 25 um.

(E) Digital karyotyping showing normal karyotype detected in SCD patient derived iPSCs (passage 4) over 8 commonly mutated regions across the genome, compared to control.

(F) HDR based correction of the sickle cell mutation in patient iPSCs using dsDNA donor shown as percentage of HDR events at the target. Error bars represent SEM (3 independent experiments). Types of HDR detected are represented below. T = Test, C = Control.
